# *SnToxA, SnTox1* and *SnTox3* originated in *Parastagonospora nodorum* in the Fertile Crescent

**DOI:** 10.1101/2020.03.11.987214

**Authors:** Fariba Ghaderi, Bahram Sharifnabi, Mohammad Javan-Nikkhah, Patrick C. Brunner, Bruce A. McDonald

## Abstract

The center of origin of the globally distributed wheat pathogen *Parastagnospora nodorum* has remained uncertain because only a small number of isolates from the Fertile Crescent, a region in the Middle East where wheat was domesticated from wild grasses, were included in earlier population genetic and phylogeographic studies. We isolated and genetically analyzed 193 *P. nodorum* strains from three naturally infected wheat fields distributed across Iran, a country located within the Fertile Crescent, using eleven neutral microsatellite loci. Compared to previous studies that included populations from North America, Europe, Africa, Australia and China, the populations from Iran had the highest genetic diversity globally and also exhibited greater population structure over smaller spatial scales, patterns typically associated with a species’ center of origin. Genes encoding the necrotrophic effectors *SnToxA, SnTox1* and *SnTox3* were found at a high frequency in the Iranian population. By sequencing 96 randomly chosen Iranian strains, we detected new alleles for all three effector genes. Analyses of allele diversity showed that all three effector genes had higher diversity in Iran than in any population included in previous studies, with Iran acting as a hub for the effector diversity that was found in other global populations. Taken together, these findings support the hypothesis that *P. nodorum* originated either within or nearby the Fertile Crescent with a genome that already encoded all three necrotrophic effectors during its emergence as a pathogen on wheat. Our findings also suggest that *P. nodorum* was the original source of the *ToxA* genes discovered in the wheat pathogens *Phaeosphaeria avenaria* f. sp. *tritici* 1, *Pyrenophora tritici-repentis* and *Bipolaris sorokiniana*.

## INTRODUCTION

*Parastagonospora nodorum* (syn. *Phaeosphaeria nodorum*) is the causal agent of Stagonospora nodorum leaf and glume blotch (SNB) on durum and bread wheat (Quaedvlieg et al. 2013). This disease is found in most wheat-growing regions of the world (Wiese, 1987) and can cause yield losses of up to 31% (Bhathal et al. 2003). The pathogen infects mainly leaves and ears, reducing both grain quality and yield (Eyal 1987, 1999).

The genetic structure of *P. nodorum* populations has been analyzed at field, regional, continental, and global scales using several types of neutral genetic markers, including restriction fragment length polymorphisms (RFLPs) (McDonald et al. 1994; Keller et al. 1997a, 1997b), amplified fragment length polymorphisms (AFLPs) (Bennett et al. 2005) and microsatellites (also called simple sequence repeats or SSRs) (Stukenbrock et al. 2005). Populations of *P. nodorum* exhibited high levels of genetic diversity in North America, Europe, Africa, Australia and China. Migration rates between continents were high, resulting in a shallow population structure even on continental and global scales (Stukenbrock et al. 2006).

Based on the findings of higher private allelic richness at eight microsatellite loci and a higher number of private multilocus haplotypes across four sequence loci in Iran compared to global populations, it was hypothesized that *P. nodorum’s* center of origin coincides with its wheat host in the Fertile Crescent (McDonald et al. 2012). However, the origin of *P. nodorum* remained uncertain because only 24 strains from the Fertile Crescent region were analyzed previously. McDonald *et al.* (2013) examined the evolutionary histories of three necrotrophic effectors (NEs) in *P. nodorum* encoded by the genes *SnToxA, SnTox1*, and *SnTox3*. Contrary to expectations, the 24 Iranian isolates did not exhibit the highest genetic diversity for any of these NEs. Instead, the highest diversity for *SnToxA* based on rarefaction analyses and the number of private alleles was observed in South Africa while the highest diversity for *SnTox1* was in Europe and the highest diversity for *SnTox3* was in North America. These findings, coupled with the absence of all three NE-encoding genes in all but one close relative of *P. nodorum* called *Phaeosphaeria avenaria* f. sp. *tritici* 1 (*Pat1*), led to the hypothesis that *P. nodorum* acquired these NEs through three independent horizontal gene transfers (McDonald et al. 2013).

Here we report new findings based on analyses of 193 new *P. nodorum* strains sampled from three naturally infected wheat fields located in different regions of Iran. We aimed to specifically test the hypothesis that the Fertile Crescent, represented by Iran, is the center of origin of *P. nodorum* by analyzing population genetic structure using the same neutral SSR loci that were used previously to analyze globally distributed *P. nodorum* populations. Measures of genotype diversity, disequilibrium and mating type frequencies were combined to determine the importance of sexual recombination in these Iranian populations. We also analyzed sequence diversity for *SnToxA, SnTox1* and *SnTox3* to test the hypothesis that all three genes were acquired by *P. nodorum* through independent horizontal gene transfers. Our findings support an origin for *P. nodorum* in the Fertile Crescent and suggest that all three necrotrophic effectors emerged in *P. nodorum* populations at their center of origin, contradicting the earlier hypothesis of independent origins through a series of horizontal gene transfers after the pathogen moved to different continents. These new data strengthen the hypothesis that *P. nodorum* was the ultimate source of the *ToxA* genes horizontally acquired by *Pat1, Pyrenophora tritici-repentis* and *Bipolaris sorokiniana*.

## MATERIALS AND METHODS

### Comparisons with earlier studies

To make comparisons with previous *P. nodorum* studies as compatible as possible, we conducted the same analyses using the same software whenever possible. To increase the overall sample size and to obtain a more detailed portrait of *P. nodorum* population structure in Iran, we included the data set of the previously described population from the northern Golestan province (McDonald et al. 2012) in our microsatellite analyses. To account for possible differences in binning of SSR alleles between the two studies, randomly chosen Golestan isolates were PCR-amplified and included in the same runs as the new Iranian isolates.

Earlier publications that analyzed the same mating types, SSR loci and necrotrophic effectors included 693 strains from 17 wheat fields located in Australia (n=73), China (n=101), Europe (n=301), Mexico (n=31), North America (n=132), and South Africa (n=55) (Stukenbrock et al. 2006), as well as 57 strains from two wheat fields sampled in 2005 and 2010 in the Golestan Province in Iran (McDonald et al, 2012). Most of these wheat fields were sampled using the same hierarchical transect sampling method that was used for the new collections from Iran reported here. Iran is considered a proxy for the Fertile Crescent because it is the only country from this region where field populations were analyzed. Among the 57 Iranian isolates analyzed earlier, more than half were shown to be closely related to *P. nodorum* (providing additional evidence for a center of origin in the Fertile Crescent, as described in McDonald et al., 2012), but 29 of these isolates were *P. nodorum*, with complete SSR and mating-type datasets obtained from the 24 Golestan strains included in the new analyses described here.

### Sampling and DNA extraction

A total of 193 new *P. nodorum* isolates were made from infected leaves and ears collected from three naturally infected wheat fields representing the major wheat-growing areas of southern Iran in the Kohgiluyeh, Khuzestan and Fars provinces (Figure 1). Including the Golestan population, these Iranian wheat fields were separated by 250 - 800 km and differed with regard to climate, wheat cultivars and wheat-growing seasons. Isolates were obtained using hierarchical sampling (McDonald et al. 1995) from six to eight spots separated by 10 m within each field. Isolates from Kogiluyeh and Khuzestan were collected from infected leaves while isolates from Fars were collected from infected ears. Only one isolate was collected from each plant. Single-spore isolation and other culturing procedures were performed as described by Halama and Lacoste (1991). Pure cultures of each isolate were stored on lyophilized filter paper strips at −80°C.

**Figure 1.**
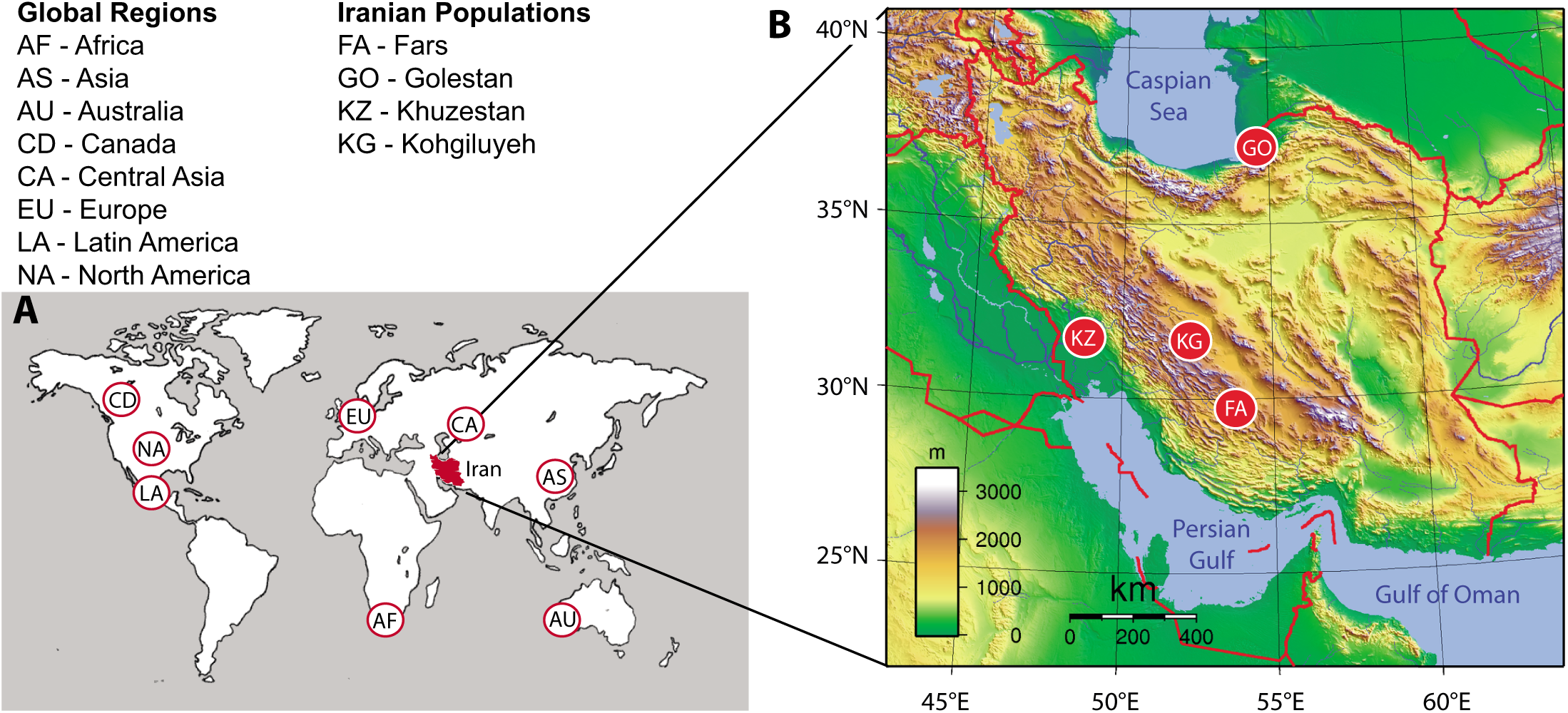
Sampling locations for *Parastagonospora nodorum* populations. **A.** The world map shows the locations of global populations described in earlier studies (Stukenbrock et al. 2006; McDonald et al. 2012). **B.** The map of Iran shows the newly sampled locations of Khuzestan, Kohgiluyeh and Fars that form the new Iranian data set as well as the earlier described population of Golestan (McDonald et al. 2012) that forms the old Iranian data set.

Isolates were grown on Petri dishes containing yeast sucrose agar (YSA, 10g/L yeast extract, 10g/L sucrose, 1.2% agar) amended with 50 μg of kanamycin. Single colonies were transferred to flasks containing 50 ml yeast sucrose broth (YSB, 10g/L yeast extract, 10g/L sucrose) and grown on an orbital shaker for 5 to 7 days at 120 rpm and 18°C. Genomic DNA was extracted as described previously (Murray and Thompson, 1980).

### Mating type determination

Mating type idiomorphs for each isolate were determined using the mating type primers described previously (Bennett et al. 2003). These primers amplified a 510 bp PCR product for *MAT1-2* isolates and a 360 bp PCR product for *MAT1-1.* Multiplex PCR amplifications were performed as described previously (Sommerhalder et al. 2006). We assessed the ratio of *MAT1-1* to *MAT1-2* alleles and tested for deviations from a 1:1 ratio of the two mating types in each field using χ ^2^ statistics.

### Microsatellite analysis

Eleven previously described SSR loci, SNOD1, SNOD3, SNOD5, SNOD8, SNOD11, SNOD15, SNOD16, SNOD17, SNOD21, SNOD22 and SNOD23 (Stukenbrock et al. 2005) were amplified in each isolate. PCR amplifications were performed in 20 μl reactions containing 0.05 μM of each primer (Microsynth, Balgach Switzerland), 1X Dream Taq Buffer (MBI Fermentas), 0.4 μM dNTPs (MBI Fermentas) and 0.5 units of Dream Taq DNA polymerase (MBI Fermentas). The PCR cycle parameters were: 2 min initial denaturation at 96°C followed by 35 cycles at 96°C for 30 sec, annealing at 56°C for 45 sec, and extension at 72°C for 1 min. A final 7 min extension was made at 72°C. Amplicons were separated in a 3730xl ABI Genetic Analyzer capillary sequencer (Life Technologies, Applied Biosystems). The software Genemapper (Life Technologies, Applied Biosystems) was used for genotyping.

### Population genetic analyses

Genetic variation was quantified using measures of gene and genotype diversity at the SSR loci. A clone-corrected data set containing a single representative of each multilocus haplotype based on the program Genodive 2.0 (Meirmans and Van Tienderen 2004) was used for subsequent analyses. Genetic diversity for each locus and for each population across all loci was assessed using the program POPGENE32 (Yeh et al. 1999). Genotype diversity and clonal fractions for all populations were calculated according to Stoddart and Taylor (1988) as implemented in the R package poppr (Kamvar et al. 2014). The percentage of maximum possible diversity (G/N) was calculated by dividing the genotypic diversity by the number of isolates (Chen et al. 1994). Allelic richness as a measure of within population genetic diversity was calculated using the program Fstat (Goudet 2001). Measures of recombination including the indices of association I_A_ and r_d_ were estimated with the program MULTILOCUS version 1.2.2 using 10^3^ simulations (Agapow and Burt 2001).

Pairwise genetic differentiation among populations was estimated as *R*_*ST*_ using ARLEQUIN 3.5 (Excoffier and Lischer 2010). In contrast to measures of *F*_*ST*_ that are based on infinite allele models, *R*_*ST*_ considers stepwise mutations to better model the evolutionary relatedness of microsatellites by counting the sum of the squared number of repeat differences between two haplotypes (Weir and Cockerham 1984; Michalakis and Excoffier 1996). P-values were obtained using 10^3^ permutations.

We analyzed the population structure of *P. nodorum* using STRUCTURE (Pritchard et al. 2000; Falush et al. 2003). The number of populations tested ranged from *K* = 1 to *K* = 7. STRUCTURE runs were performed using the admixture model, sampling locations as priors, 10^6^ iterations and a burn-in period of 30,000. Ten independent simulations were performed. CLUMPAK (Kopelman et al. 2015) was used for summation and graphical representation of the STRUCTURE runs. The implemented deltaK approach of Evanno *et al.* (2005) was used to identify the optimal number of populations in the data set.

### Sequence analyses of necrotrophic effectors

All strains were screened for the presence of *SnToxA, SnTox1*, and *SnTox3* using PCR assays. PCR amplifications were carried out in 20 μL reaction mixtures including 0.04 μM of each primer (Microsynth, Balgach Switzerland), 1X Dream Taq Buffer (MBI Fermentas), 0.4μM dNTPs (MBI Fermentas) and 0.4 units of Dream Taq DNA polymerase (MBI Fermentas). The PCR cycle parameters were: initial denaturation of 2 min at 96°C, 36 cycles of denaturation at 96°C for 40 sec, annealing temperature specific for each primer for 45 sec, extension at 72°C for 1 min followed by a final extension at 72°C for 7 min. PCR products were purified to remove unincorporated nucleotides and primers using NucleoFast® 96 PCR plates (Macherey-Nagel, Oensingen, Switzerland). Details of the annealing temperatures and primers were published previously (Friesen et al. 2006 for *SnToxA*; Liu et al. 2009 for *SnTox3*). The *SnTox1* primers were newly designed in this study. Sequences and annealing temperatures specific for each primer pair are listed in Supplementary Table S1.

Sequencing reactions were produced in both directions using the BigDye® Terminator v3.1 Sequencing Standard Kit (Life Technologies, Applied Biosystems). The PCR cycle parameters were 2 min at 96°C followed by 55 or 99 cycles for 10 sec at 96°C, for 5 sec at 50°C and for 4 min at 60°C. PCR products were cleaned with the Illustra Sephadex G-50 fine DNA grade column (GE Healthcare) according to the manufacturer’s recommendations and sequenced with a 3730xl Genetic Analyzer (Life Technologies, Applied Biosystems). Forward and reverse sequences were aligned using the program Sequencer 5.1 (Gene Code, Ann Arbor, MI). Final alignments were performed using MAFFT ver.7 (Katoh and Standley 2013).

## RESULTS

### Sources of isolates

We obtained new data from 193 *P. nodorum* isolates sampled from wheat fields located in three major provinces (Kohgiluyeh, Fars and Khuzestan) in the south of Iran (Figure 1; Table 1). We will refer to these isolates collectively as the “new Iranian” population. We added data from 24 isolates originating from the Golestan province in northern Iran that were described in earlier studies (McDonald et al. 2012; McDonald et al. 2013) and will refer to these isolates collectively as the “old Iranian” population. We will refer to the combination of the new and old Iranian populations as the “combined Iranian” population. The combined Iranian population came from four provinces, separated by mountains and deserts, that are characterized by different climates, cropping systems, wheat cultivars and growing seasons, with distances among Iranian field populations ranging from 250-800 km. Measures of genetic diversity in the combined Iranian populations of *P. nodorum* were compared with identical measures made in an earlier analysis that included 693 *P. nodorum* isolates sampled from 17 wheat fields coming from nine regions distributed across five continents, with none of these earlier collections coming from the Fertile Crescent (Stukenbrock et al. 2006). We will refer to these 693 isolates as the “global populations”.

**Table 1.** Collection sites, sample sizes and host source for *Parastagonospora nodorum* isolates from Iran. The population from Golestan was described in earlier studies (McDonald *et al.* 20012; McDonald et al., 2013).

| Region | Year collected | Collectors | Sample size | Host source | Climate |
| --- | --- | --- | --- | --- | --- |
| Kohgiluyeh | 2015 | F. Ghaderi | 64 | wheat leaves | temperate, mild winters |
| Fars | 2015 | F. Ghaderi | 66 | wheat ears | temperate, mild winters |
| Khuzestan | 2015 | F. Ghaderi | 63 | wheat leaves | warm, dry |
| Golestan | 2005, 2010 | R. Sommerhalder/M. Razavi | 24 | wheat ears | warm, humid |

### Measures of genetic diversity

Diversity parameters for the individual SSR loci are summarized in Supplementary Table S2. All eleven SSR loci were successfully amplified for all *P. nodorum* isolates in the new Iranian population. All loci were polymorphic, with the number of alleles ranging from 3 to 22. Gene diversity ranged from 0.18 (SNOD15) to 0.94 (SNOD1) with an average across all loci of 0.60.

Overall levels of gene and genotype diversity were high in each Iranian field population (Table 2). A total of 213 different multilocus genotypes were detected among the 217 combined Iranian isolates, corresponding to an overall clonal fraction of only 2%, with no field population showing a clonal fraction higher than 3%. By way of comparison, clonal fractions among the global populations averaged 6% and ranged from 2% in Switzerland to 33% in Mexico (Stukenbrock et al. 2006). Nei’s measure of gene diversity for the combined Iranian data set ranged from 0.56 in Golestan to 0.66 in Fars and averaged 0.60 across the combined Iranian field populations. Gene diversity among the global populations was generally lower, ranging from 0.44 in Mexico to 0.57 in Texas with an average of 0.58 when including all 693 isolates in the global population (Stukenbrock et al. 2006). Allelic richness among the combined Iranian populations was estimated by resampling 10^3^ datasets from each field population. The allelic richness varied between 4.61 in Golestan to 5.72 in Khuzestan, with an average allelic richness across the combined Iranian population of 4.85 (Table 2). We compared the allelic richness and private allelic richness at SSR loci with the global populations using rarefaction to account for differences in sample size. In agreement with a previous study that included only the old Iranian population from Golestan (McDonald et al. 2012), private allelic richness in the combined Iranian population was higher than other regions around the world. Adding the new Iranian population to the old Iranian population also led to an increase in mean allelic richness, giving Iran the highest level of neutral gene diversity on a global scale (Figure 2).

**Table 2.** Measures of gene and genotypic diversity and estimates of linkage disequilibrium based on eleven microsatellite loci and mating type ratios in the Iranian *Parastagonospora nodorum* populations

| Population | no. of isolates | Clone corrected | Clonal fraction | Genotype diversity | $I_A$ | $r_d$ | Gene diversity | Allelic richness | Ratio *MAT1-1:MAT1-2* | $\chi^2$ |
| --- | --- | --- | --- | --- | --- | --- | --- | --- | --- | --- |
| Kohgiluyeh | 64 | 62 | 0.03 | 97% | 0.31*<br>(0.21*) | 0.032*<br>(0.028*) | 0.58 | 4.99 | 45:17 | 6.66* |
| Khuzestan | 63 | 61 | 0.03 | 97% | 0.068<sup>ns</sup> | 0.008<sup>ns</sup> | 0.57 | 4.72 | 34:27 | 0.40<sup>ns</sup> |
| Fars | 66 | 66 | 0 | 100% | 0.23*<br>(0.12<sup>ns</sup>) | 0.025*<br>(0.015<sup>ns</sup>) | 0.66 | 5.10 | 37:29 | 0.49<sup>ns</sup> |
| Golestan | 24 | 24 | 0 | 100% | 0.034<sup>ns</sup> | 0.004<sup>ns</sup> | 0.56 | 4.61 | 10:14 | 0.34<sup>ns</sup> |
| All populations | 217 | 213 | 0.02 | 98% | 0.34*<br>(0.24<sup>ns</sup>) | 0.033*<br>(0.027<sup>ns</sup>) | 0.60 | 4.85 | 126:87 | 3.59<sup>ns</sup> |

$I_A$ , $r_d$ ; Association indices (measures of linkage disequilibrium). Population estimates in brackets are after removing the linked locus SNOD8. The "All populations" estimates in brackets are based on excluding the admixed population Kohgiluyeh.
\* Significant at $P < 0.05$ ; ns: non-significant.
$\chi^2$ Value for deviation from a 1:1 mating type ratio.

**Figure 2.**
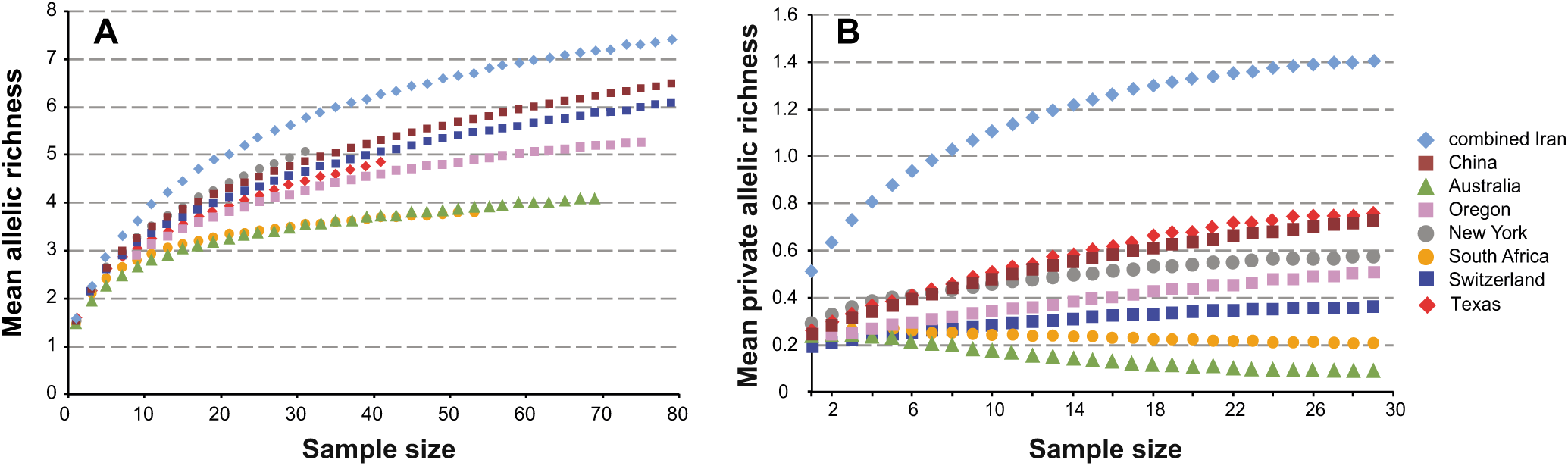
Comparisons of population diversity at neutral SSR loci for global *Parastagonospora nodorum* populations. ADZE rarefaction results for the combined Iranian populations of *P. nodorum* compared to global populations analyzed in an earlier study (McDonald et al. 2012). The rarefied mean allelic richness (**A**) and private allelic richness (**B**) are shown in relation to sample size. Maximum samples sizes are limited by the smallest population sample size.

### Population differentiation

Genetic differentiation between all Iranian populations was estimated using the stepwise R_ST_ mutation model (Supplementary Table S3). Differentiation between southern population pairs ranged from R_ST_ = 0.08 to 0.11, while differentiation between the southern populations and the northern population of Golestan was higher, ranging from R_ST_ = 0.26 to 0.37. These values were considered to be high given the much smaller geographic sampling scale compared to the nine global populations located on different continents and separated by thousands of km that showed an average pairwise R_ST_ = 0.07 for the same SSR loci (Stukenbrock et al. 2006).

In concordance with the R_ST_ analyses, STRUCTURE clustered the Iranian isolates into four distinct groups according to their geographical origins (Figure 3). The high assignment probabilities for all individuals from Golestan, Fars and Khuzestan suggest limited gene flow among these populations. In contrast, several isolates from Kohgiluyeh were identified as being admixed mainly with the Fars population, suggesting recent gene flow between these two populations.

**Figure 3.**
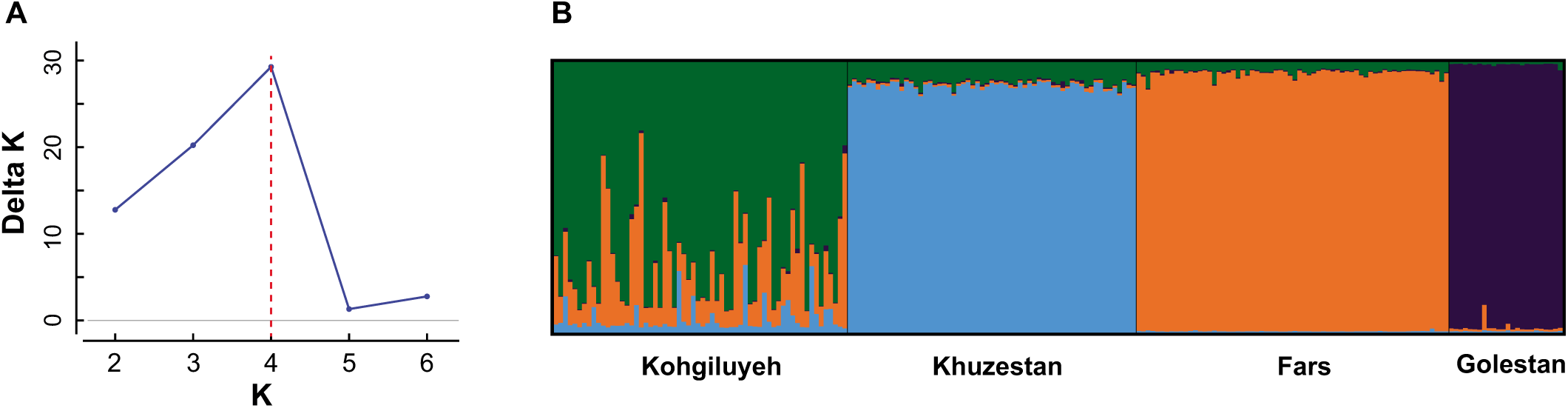
Structure analysis of the combined Iranian *Parastagonospora nodorum* populations. Individuals were assigned to clusters based on the SSR data set. **A.** DeltaK shows the highest likelihood at K = 4 assumed populations. **B.** STRUCTURE plot of assignment probabilities for each individual assuming K = 4 populations.

### Tests for random mating based on mating type frequencies and measures of linkage disequilibrium

PCR amplification of the mating type idiomorphs produced single amplicons corresponding to either *MAT1-1* or *MAT1-2* in all Iranian isolates. The frequency distribution showed a bias towards *MAT1-1*, but it was not significantly different from the expected 1:1 ratio for the entire data set, with 126 *MAT1-1* vs. 87 *MAT1-2*. The mating type ratio was not significantly different from 1:1 for three of the field populations, but Kohgiluyeh had a significantly skewed distribution with 45 *MAT1-1 vs.* 17 *MAT1-2* (Table 2).

Linkage disequilibrium (LD) estimates were significantly different from the distribution expected under the hypothesis of no associations for Kogiluyeh and Fars (Table 2). Closer inspection revealed that loci SNOD8 and SNOD17 were significantly associated. A BLAST search on the genome of the *P. nodorum* reference isolate SN15 v2.0 (Hane et al. 2007) revealed that both loci were located on scaffold 4 approximately 1000 bp apart, i.e. they were tightly linked. Removing locus SNOD8 from the analysis resulted in non-significant LD for Fars, but Kogiluyeh still showed significant LD.

### Analyses of necrotrophic effectors

We PCR-screened the new Iranian population for the presence of the *SnToxA, SnTox1* and *SnTox3* genes with gene-specific primers (Supplementary Table S1). *SnToxA* and *SnTox1* were found at frequencies of 95% and 97%, respectively while *SnTox3* was found in 72% of isolates. The distribution of all possible multi-effector genotypes is shown in Supplementary Fig. S1. The combination with all three effectors present (A+3+1+) was by far the most abundant (70%, compared to 66% expected under neutral associations), followed by genotype A+3-1+ (25%, compared to 24% expected under neutral associations). When compared to an earlier analysis of multi-effector genotypes in the global population of *P. nodorum*, only the Australian population had a higher frequency of strains carrying all three necrotrophic effectors (McDonald et al. 2013).

We sequenced *SnToxA, SnTox1* and *SnTox3* for 96 randomly chosen isolates from the new Iranian population and added the previously published sequence data from the old Iranian population (McDonald et al. 2013). The combined Iranian *SnToxA* sequences collapsed into 16 distinct haplotypes. Twelve of these haplotypes were already detected among the *P. nodorum* isolates in the global data set, and haplotypes H1, H5 and H15 had been detected in the related species *Pat1* (McDonald et al. 2013). Importantly, we detected several *SnToxA* haplotypes among the new Iranian isolates that were previously reported as private alleles for other regions (McDonald et al. 2013). For example, *SnToxA* haplotypes H2, H4 and H9 were unique to South Africa in the earlier global data set, but we found each of these alleles in the new Iranian field populations (Figure 4). Three *SnToxA* haplotypes (H18 – H20) found in the new Iranian population were not detected previously, resulting in Iran having the highest number of private *SnToxA* alleles and increasing the number of distinct *P. nodorum SnToxA* haplotypes to 20 (Figure 4A). The 20 *SnToxA* haplotypes translated into 11 protein isoforms. It is noteworthy that all 11 isoforms were detected among Iranian isolates (Figure 4B). Haplotypes 3 and 17, detected in North America and South Africa, respectively, contain several stop codons each and are unlikely to translate into functional proteins (Figure 4).

**Figure 4.**
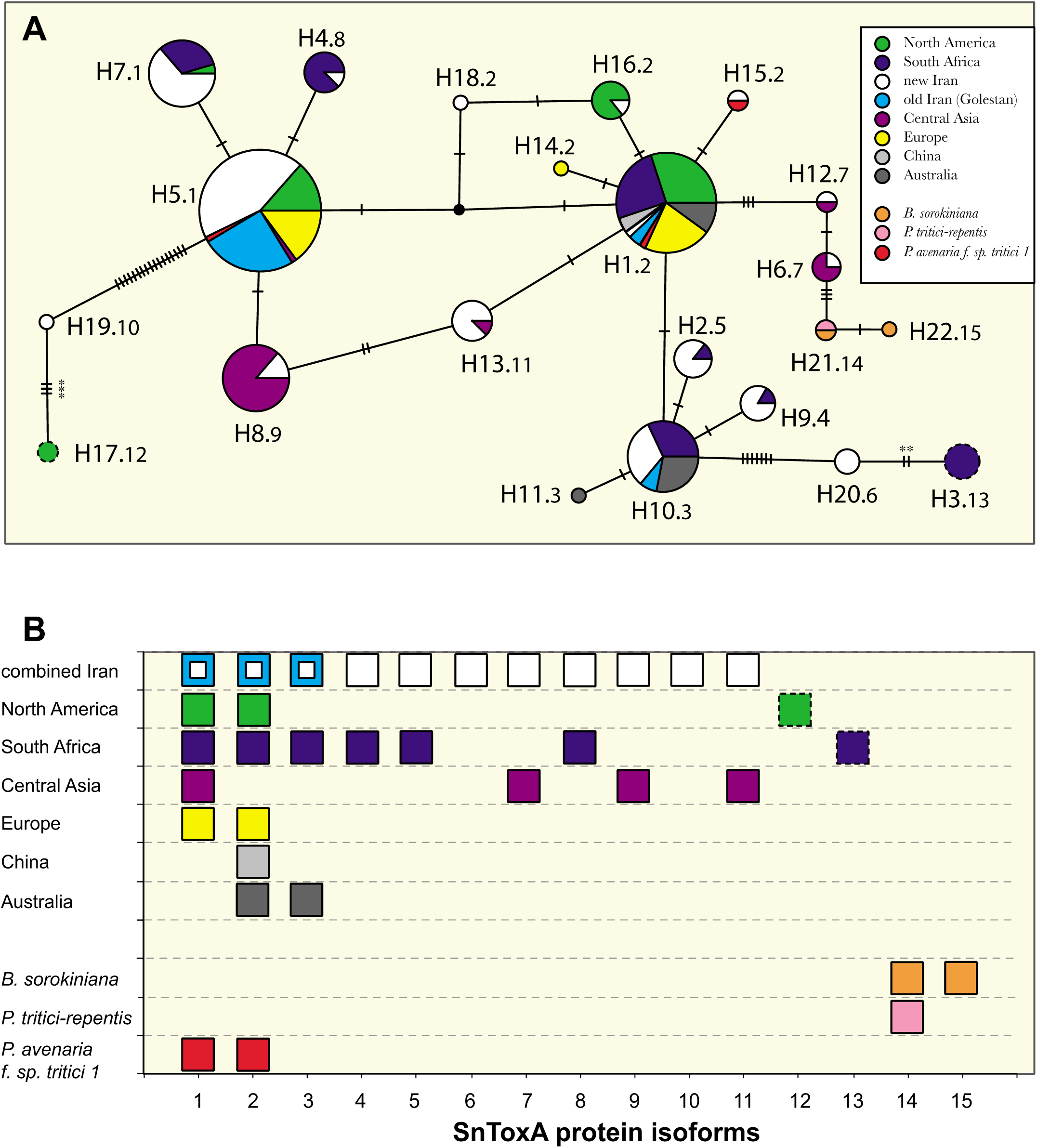
Haplotype network showing global *SnToxA* diversity. **A.** A TCS network connecting all distinct *SnToxA* haplotypes. Populations organized according to global regions use the same color-coding as McDonald et al. (2013) to enable easy comparisons. The new Iranian isolates (in white) are indicated separately from the old Iranian isolates (in blue). The sizes of haplotype circles are proportional to their frequencies in the *P. nodorum* data set. Haplotypes found in other species than *P. nodorum* are represented by a single individual. The number after the dot in the haplotype designation indicates the corresponding protein isoform. Hash marks indicate single nucleotide polymorphisms and asterisks indicate stop codons. **B.** The distribution of SnToxA protein isoforms across populations. Isoforms 12 and 13 (surrounded by dashed lines) are likely to be non-functional because of several stop codons in both nucleotide haplotypes.

Similar patterns of diversity were observed for *SnTox1* and *SnTox3*, leading the combined Iranian population to have the highest number of private alleles for all three NE-encoding genes. The combined Iranian *Tox1* sequences collapsed into 11 distinct haplotypes. Of these, four haplotypes (H19 - H22) were new and unique to Iran, increasing the total number of *SnTox1* haplotypes to 22 (Supplementary Figure S2). The combined Iranian *SnTox3* sequences collapsed into eight distinct haplotypes. Of these, two haplotypes were new and unique to Iran (H12 – H13), increasing the total number of *SnTox3* haplotypes to 13 (Supplementary Figure S3). The rarefaction analyses also identified Iran as the region with the highest number of *SnToxA* alleles (Figure 5), but other regions had higher numbers of *SnTox1* and *SnTox3* alleles (Supplementary Figure S4).

**Figure 5.**
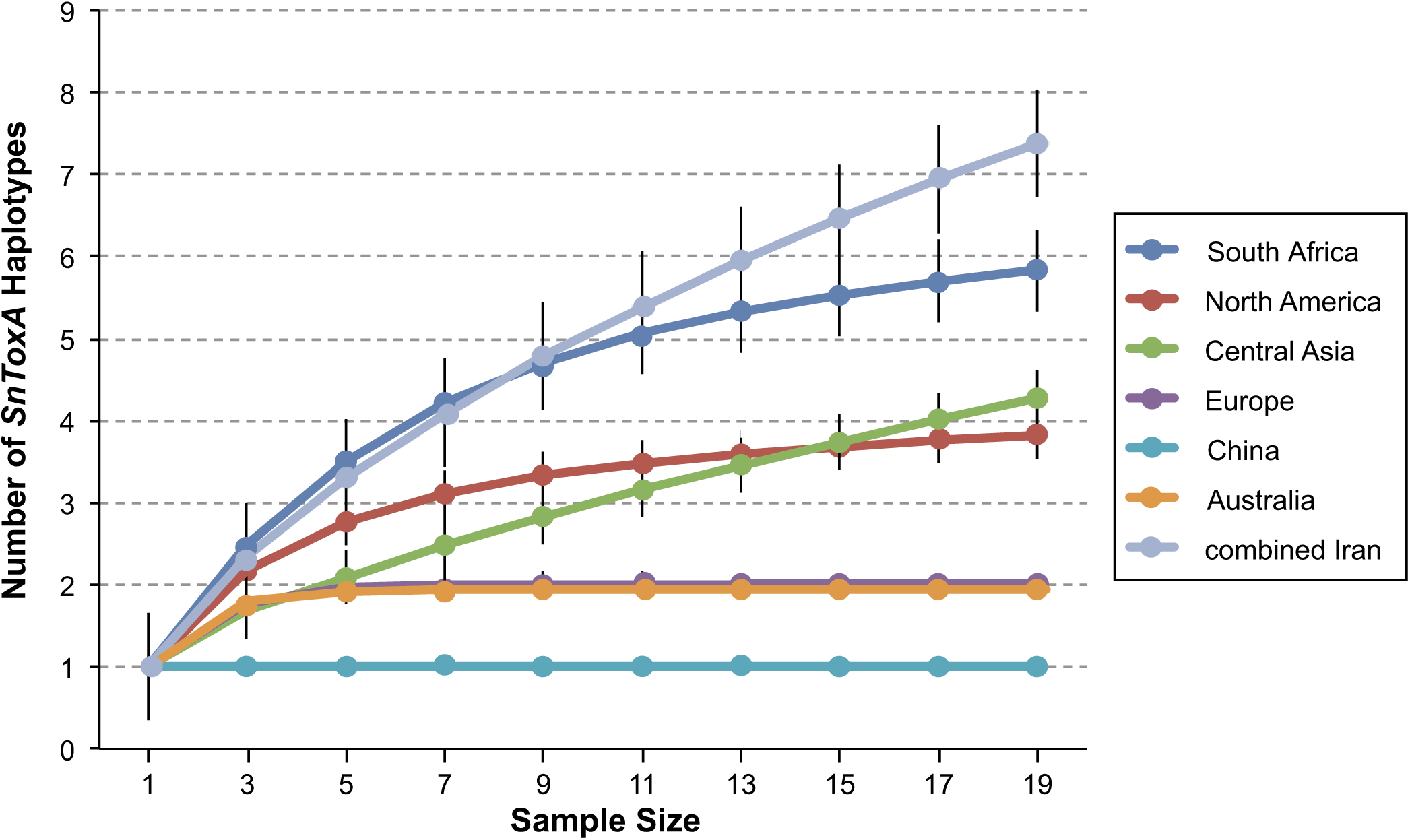
Rarefaction analysis of global *SnToxA* diversity. The rarefied number of *SnToxA* alleles found in each resampled population as a function of increasing sample size. Data for non-Iranian populations were taken from an earlier study (McDonald et al. 2013).

Because our sequence analyses detected many Iranian effector alleles that were shared with global populations (Figure 4, Supplementary Figures S2, S3), we conducted a more detailed analysis of the number of effector alleles that were shared between regional populations. Iran shared the highest number of effector alleles with other populations for *SnToxA* and *SnTox3*, but the pattern was less clear for *SnTox1* because several populations had similar numbers of shared alleles (Supplementary Fig. S5).

## DISCUSSION

We analyzed population genetic diversity for neutral and selected loci in a large collection of *P. nodorum* isolates obtained from four different regions in Iran. We found that the Iranian field populations had the highest genetic diversity detected globally to date for all genetic markers. We also found significant population structure in Iran using neutral SSR markers. These results add strong support to the hypothesis that *P. nodorum* originated in the same region where wheat was domesticated (Balter 2007), similar to what was found for the wheat pathogen *Zymoseptoria tritici* (Banke and McDonald 2005; Stukenbrock et al. 2006). We also discovered that Iran is a global hotspot of diversity for all three genes encoding necrotrophic effectors and that Iran appears to be a hub or source population for effector alleles shared with other *P. nodorum* populations around the world. Taken together, these findings support the hypothesis that the Fertile Crescent population of *P. nodorum* is the source population for all three effectors, arguing against an earlier hypothesis that all three effectors were acquired by *P. nodorum* through separate horizontal transfers after it escaped from the Fertile Crescent (McDonald et al. 2013).

### *P. nodorum* originated in the Fertile Crescent

It is generally assumed that populations at a species’ center of origin (i.e. the original source for other populations that became established in new locations) will display the highest genetic diversity at neutral loci due to the accumulation of new mutations over many generations. In contrast, more recently founded sink populations typically show lower genetic diversity compared to older source populations as a result of genetic bottlenecks imposed by the founding event coupled with less residence time for mutations to accumulate.

Consistent with a center of origin, the Iranian populations of *P. nodorum* showed the highest diversity at neutral SSR loci compared to other global populations. The average allelic richness across the combined Iranian populations was 4.85, while the highest allelic richness found in global populations was 4.44 (Switzerland) and the mean allelic richness based on combining all 693 global isolates was 4.81 (Stukenbrock et al. 2006). Rarefaction analyses also showed that Iran has the highest allelic richness for SSRs compared to all other global populations (Fig. 2).

Despite the relatively small spatial scale sampled in Iran, the four Iranian populations showed strong signatures of differentiation, with an average R_ST_ of 0.20. For comparison, the differentiation between global populations separated by similar distances were R_ST_ = 0.01 between Texas and Oregon, R_ST_ = 0.00 between Denmark and Switzerland, and the average differentiation among all global populations was 0.07 (Stukenbrock et al. 2006). The finding of greater genetic structure among geographically nearby populations is expected for older populations found at a species’ center of origin as a result of mutation/drift equilibrium reached after many generations among populations separated by geographical features such as mountains and deserts (see review by Orsini et al. 2013 and references therein).

Consistent with this theory, the STRUCTURE analyses showed that the four Iranian sampling sites maintained genetically distinct *P. nodorum* populations. While Golestan, Fars and Khuzestan showed only marginal signatures of admixture, several isolates from Kohgiluyeh were identified as being admixed with Fars (Fig. 3). We speculate that this signature of gene flow is a result of human-mediated dispersal of infected seed from the Fars region into the Kohgiluyeh region, which enabled introgression of *P. nodorum* alleles from the Fars population into the population at Kohgiluyeh. *P. nodorum* often infects ears and seed-borne infection is common (e.g. Bennett et al. 2007). Infected seeds are thought to be the main mechanism responsible for the global dispersal of the pathogen (Bennett et al. 2005).

Regular sexual recombination is a key factor that affects a pathogen’s evolutionary potential. The reshuffling of polymorphism allows for rapid adaptation both to the host’s immune response and to external stresses such as the application of fungicides (McDonald and Linde 2002; Linde et al. 2003). In previous studies, most populations of *P. nodorum* were reported to exhibit the “signature of sex”, including high genotypic diversity, low clonality, random associations among neutral markers (i.e. gametic equilibrium), and approximately equal frequencies of the two mating types (Bennet et al. 2005; Stukenbrock et al. 2006). Our findings in Iran were largely in agreement with these earlier studies. After removing one of two linked microsatellite markers, three of the Iranian populations were in gametic equilibrium, but the Kohgiluyeh population showed gametic disequilibrium and mating type frequencies that deviated significantly from the expected 1:1 ratio. We also found that many strains from Kohgiluyeh showed strong signatures of population admixture, mainly with Fars. Admixture can result in high levels of LD that will decline over time due to recombination (Pritchard and Rosenberg 1999). Based on our findings, we hypothesize that the skewed mating type ratio and linkage disequilibrium in Kohgiluyeh are a result of recent admixture with *P. nodorum* isolates from the Fars region following a recent introduction on infected seed.

### Evidence that all three necrotrophic effectors originated in *P. nodorum* at its center of origin

*P. nodorum* possesses several necrotrophic effectors that facilitate the infection process (Friesen et al. 2008). These NEs show population-specific patterns of presence/absence polymorphism as well as high nucleotide diversity consistent with local adaptation to the NE sensitivity genes carried by the local host population (Stukenbrock and McDonald 2007; McDonald et al. 2013). Earlier analyses of nucleotide diversity in the NE-encoding genes *SnToxA, SnTox1* and *SnTox3* did not provide evidence for a single center of origin for these effectors (Stukenbrock and McDonald 2007; McDonald et al. 2013). The highest nucleotide diversity for *SnToxA* was found in South Africa, the highest diversity for *SnTox1* was in Europe, and the highest diversity for *SnTox3* was in North America. These findings, coupled with the discovery that all three effector genes were found only in the closely-related *Parastagonospora* lineages *P. nodorum* and *Pat1* and not in more distant *Parastagonospora* lineages (McDonald et al. 2012; McDonald et al. 2013), led to the hypothesis that *P. nodorum* acquired all three NEs through independent horizontal gene transfers (McDonald et al. 2013). Our detailed analyses presented here, that included a much larger and more geographically diverse collection of strains from Iran, indicate a very different scenario, namely an origin for all three NEs in the *P. nodorum* population residing in the Fertile Crescent. We detected the highest number of private alleles for all three NEs in the Iranian populations and we discovered that many alleles previously reported to be unique in other global populations also existed in Iran. For example, earlier analyses indicated that South Africa had four private alleles for *SnToxA* (haplotypes H2, H3, H4 and H9). Apart from H9, all of these alleles were found in the new *P. nodorum* collections from Iran (Fig. 4A).

It is noteworthy that many effector alleles were shared between very distant populations. For example, the most frequent allele for *SnToxA* (H1) was found in Iran, North America, Australia, Europe and also in South Africa. Some very rare alleles were also shared among distant populations. For example, *SnToxA* alleles H2 and H9 were previously found only in single isolates from South Africa (McDonald et al. 2013). In our new Iranian population, we found 6 isolates with the H2 allele and 5 isolates with the H9 allele. This finding of shared rare alleles can be interpreted as evidence for recent gene flow between Iran and South Africa. An alternative explanation is that the same allele emerged independently in the two populations, perhaps as a result of selection due to deployment of the same effector sensitivity genes in the corresponding host populations. To investigate the pattern of shared alleles in more detail, we conducted pairwise population analyses of normalized numbers of shared effector haplotypes (Supplementary Fig. S5). Iran was identified as the region that shared the most haplotypes with all other populations for *SnToxA* and *SnTox3*. The pattern for *SnTox1* was inconclusive because several populations shared similar numbers of haplotypes.

We conclude that the observed patterns of SSR diversity and necrotrophic effector diversity are most compatible with the hypothesis that *P. nodorum* originated in the Fertile Crescent region and that the original pathogen populations already carried the genes encoding *SnToxA, SnTox1* and *SnTox3*. More population samples from other regions in the Fertile Crescent, eg. Turkey, Israel or Iraq, should be analyzed to further test this hypothesis. Our results do not support an earlier hypothesis of independent origins for the three necrotrophic effectors through horizontal gene transfer. The *ToxA* gene was also discovered in the genomes of the wheat tan spot pathogen *Pyrenophora tritici-repentis* (Friesen et al. 2006) and the wheat spot blotch pathogen *Bipolaris sorokiniana* (McDonald et al. 2018), though only one allele (H21 in Figure 4A) has been found in *P. tritici-repentis* and only two alleles (H21 and H22 in Figure 4A) have been found in *B. sorokiniana*. Though H21 and H22 have not yet been found in *P. nodorum*, the closest allele in *P. nodorum* (H6) was found in Iran and is separated from H21 by only three SNPs. Taken together, these findings support the hypothesis that *P. nodorum* was the original source of the *ToxA* genes now found in *Pat1* (McDonald et al. 2013), *Pyrenophora tritici-repentis* (Friesen et al. 2006) and *Bipolaris sorokiniana* (McDonald et al. 2018).

## ACKNOWLEDGMENTS

The authors are grateful to M. Zala for his assistance in the laboratory. DNA data were collected in the Genetic Diversity Centre of ETH Zurich. FG was supported by a PhD scholarship from the Iranian Ministry of Higher Education. This work was funded by the Swiss Federal Institute of Technology (ETH), Zurich.

## Appendix: Supplementary Tables

**Table S1.**
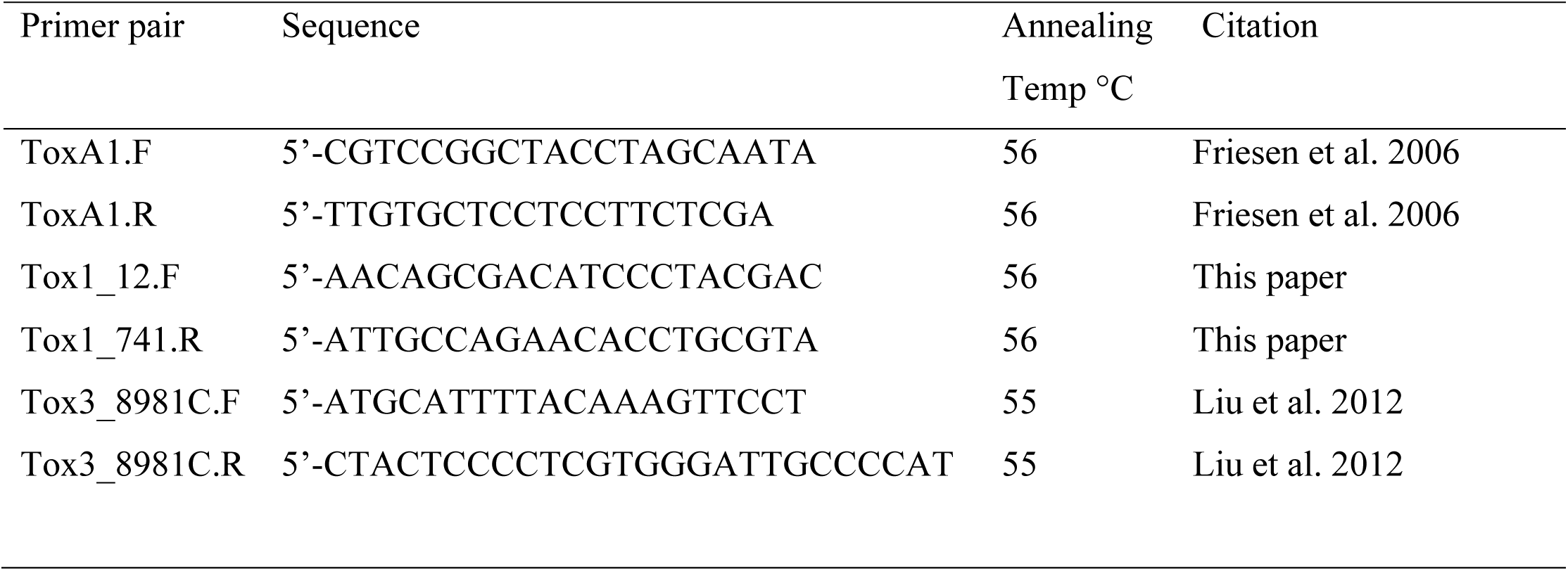
PCR primers used for the detection of the necrotrophic effectors *SnToxA, SnTox1*, and *SnTox3* among *P. nodorum* isolates.

**Table S2.** Measures of microsatellite diversity for Iranian samples of *Parastagonospora nodorum*

| Locus | Size range | Fars<br>(N = 66) | Kohgiluyeh<br>(N = 64) | Khuzestan<br>(N = 63) | Golestan<br>(N = 23) | Total no. of<br>alleles | Gene<br>diversity<sup>†</sup> |
| --- | --- | --- | --- | --- | --- | --- | --- |
| SNOD1 | 253-439 | 13 | 14 | 20 | 13 | 22 | 0.94 |
| SNOD3 | 278-318 | 3 | 3 | 2 | 2 | 4 | 0.28 |
| SNOD5 | 409-472 | 12 | 8 | 8 | 5 | 16 | 0.88 |
| SNOD8 | 325-425 | 7 | 7 | 1 | 4 | 10 | 0.71 |
| SNOD11 | 231-236 | 3 | 2 | 2 | 2 | 4 | 0.5 |
| SNOD15 | 156-174 | 2 | 2 | 2 | 3 | 3 | 0.18 |
| SNOD16 | 188-208 | 5 | 5 | 6 | 6 | 9 | 0.5 |
| SNOD17 | 92-158 | 3 | 4 | 7 | 5 | 10 | 0.68 |
| SNOD21 | 191-228 | 7 | 6 | 5 | 5 | 7 | 0.79 |
| SNOD22 | 225-267 | 6 | 6 | 4 | 5 | 8 | 0.73 |
| SNOD23 | 294-402 | 4 | 7 | 4 | 1 | 7 | 0.33 |

<sup>†</sup>Calculated according to Nei (1973)

**Table S3.** Population differentiation measured by pairwise R_ST_ (above diagonal) and corresponding p*-*values (below diagonal) among the four Iranian *Parastagonospora nodorum* populations.

| Population | Kohgiluyeh | Khuzestan | Fars | Golestan |
| --- | --- | --- | --- | --- |
| Kohgiluyeh | --- | 0.076 | 0.042 | 0.362 |
| Khuzestan | 0.005 | --- | 0.114 | 0.257 |
| Fars | 0.014 | < 0.001 | --- | 0.368 |
| Golestan | < 0.001 | < 0.001 | < 0.001 | --- |

## Appendix: Supplementary Figures

**Figure S1.**
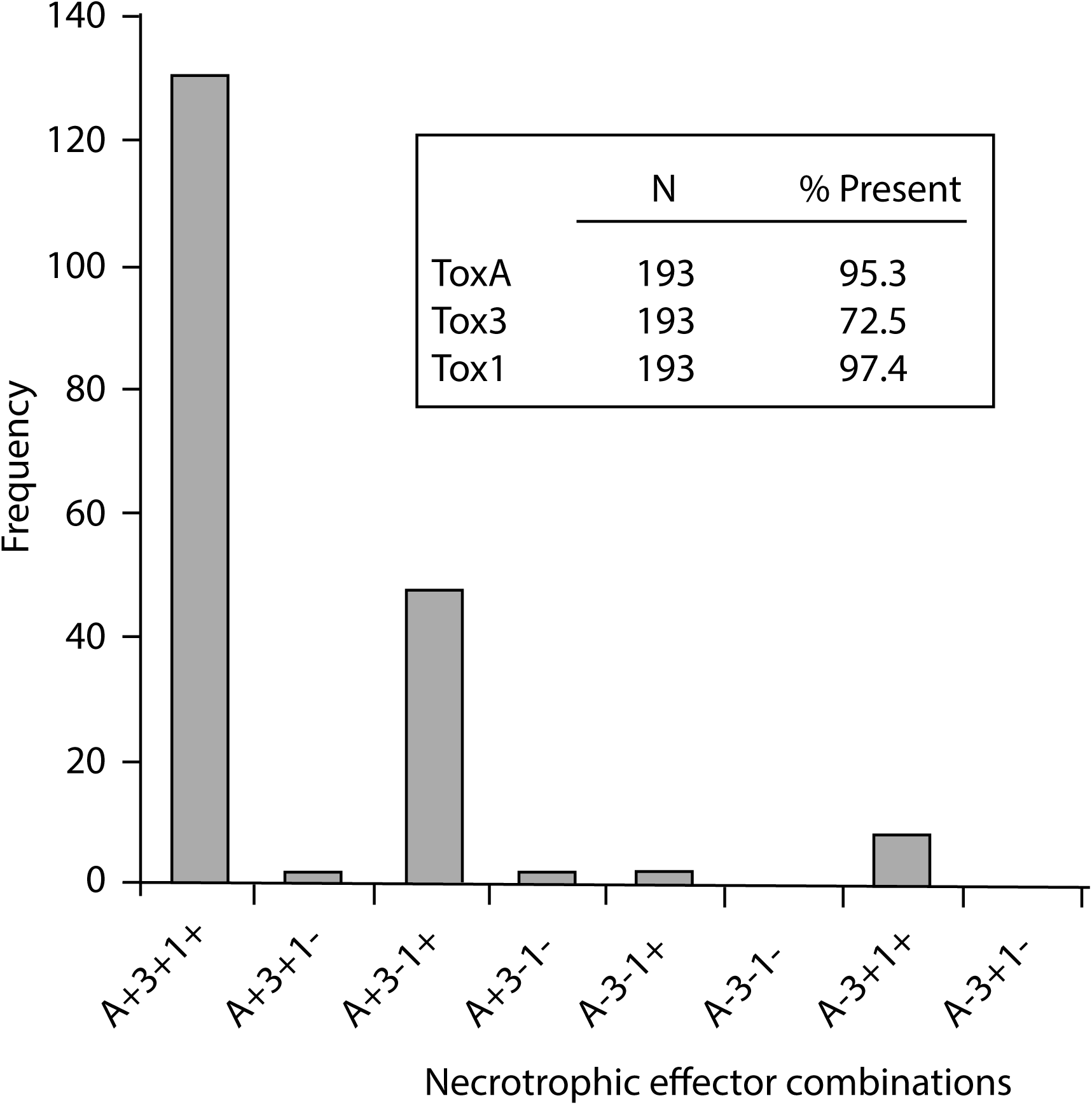
Frequency distribution of multi-effector genotypes in the new Iranian population of *Parastagonospora nodorum*. Each isolate was screened for the presence of *SnToxA, SnTox1* and/or *SnTox3* using gene-specific PCR primers. The X-axis shows all possible combinations of presence (+) and absence (-) for each effector. The legend shows the total number of individuals assayed and the percentage carrying each effector.

**Figure S2.**
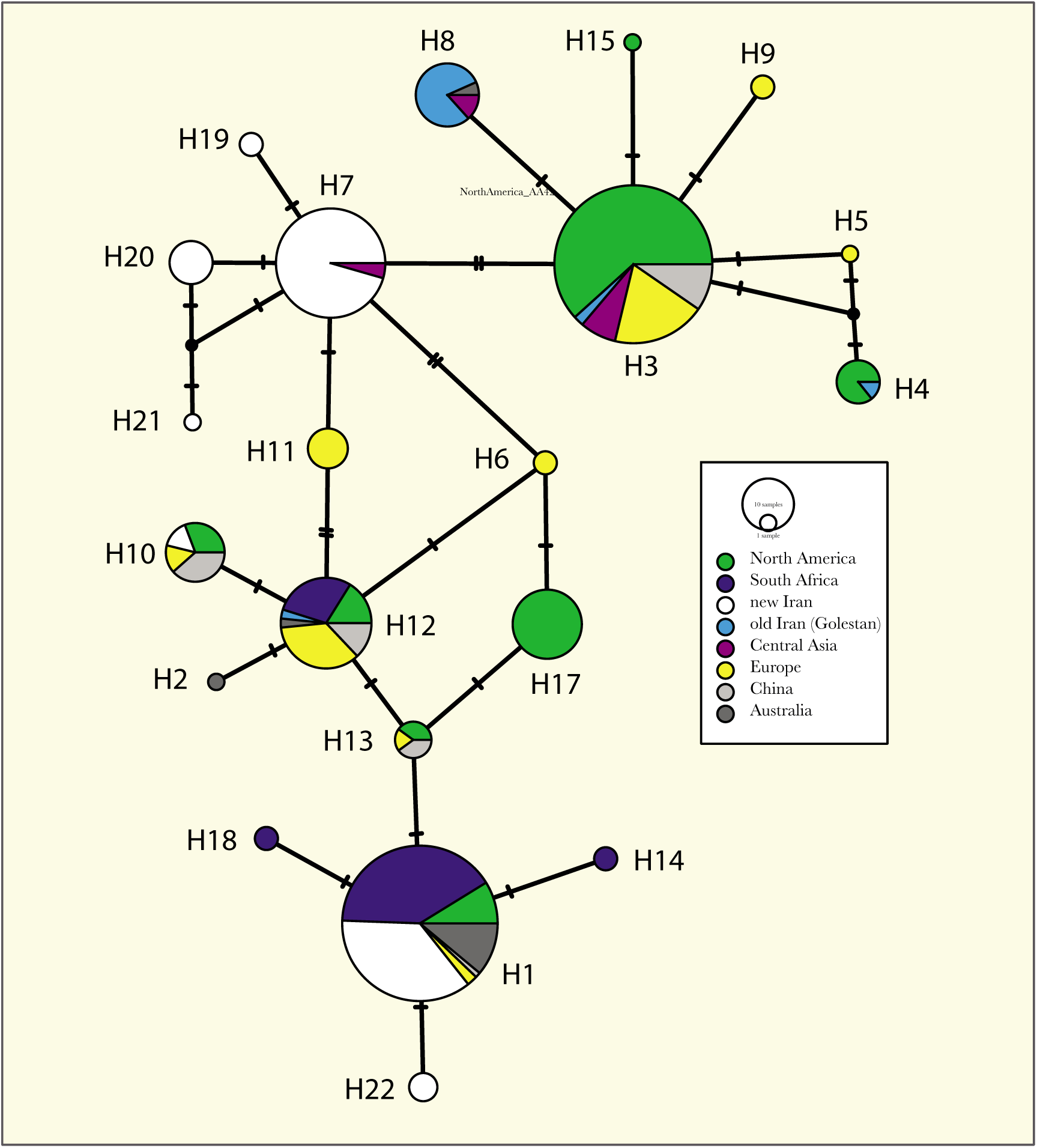
Haplotype network showing global *SnTox1* diversity. The TCS network connects all distinct *SnTox1* haplotypes. Populations organized according to global regions use the same color-coding as McDonald *et al.* (2013) to enable easy comparisons. The new Iranian isolates (in white) are indicated separately from the old Iranian isolates (in blue). The sizes of haplotype circles are proportional to their frequencies. Hash marks indicate single nucleotide polymorphisms.

**Figure S3.**
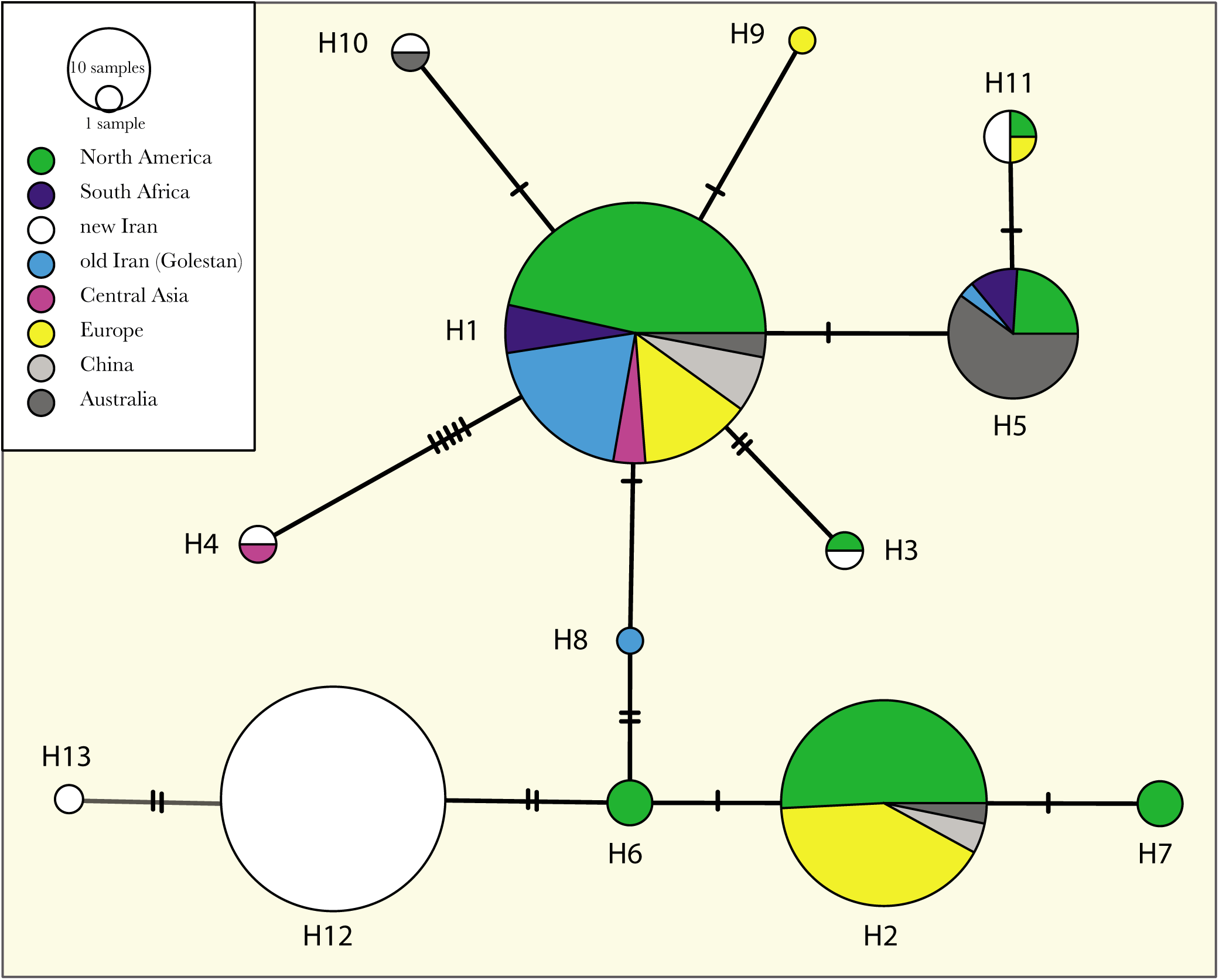
Haplotype network showing global *SnTox3* diversity. The TCS network connects all distinct *SnTox3* haplotypes. Populations organized according to global regions use the same color-coding as McDonald *et al.* (2013) to enable easy comparisons. The new Iranian isolates (in white) are indicated separately from the old Iranian isolates (in blue). The sizes of haplotype circles are proportional to their frequencies. Hash marks indicate single nucleotide polymorphisms.

**Figure S4.**
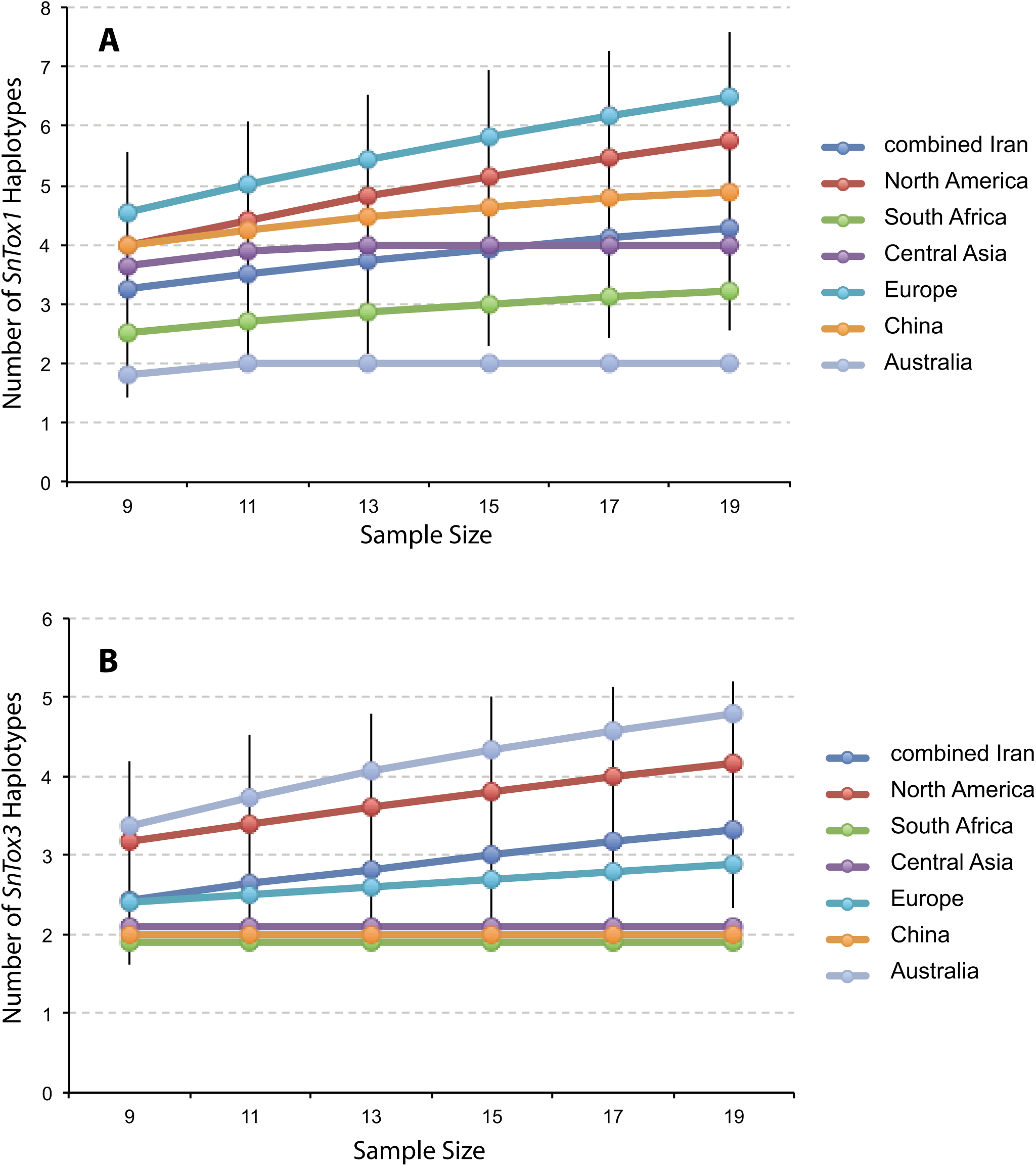
Rarefaction analysis of global *SnToxA* diversity. **A**. The rarefied number of *SnTox1* alleles. **B**. The rarefied number of *SnTox3* alleles found in each resampled population as a function of increasing sample size. Data for non-Iranian populations were taken from an earlier study (McDonald et al. 2013).

**Figure S5.**
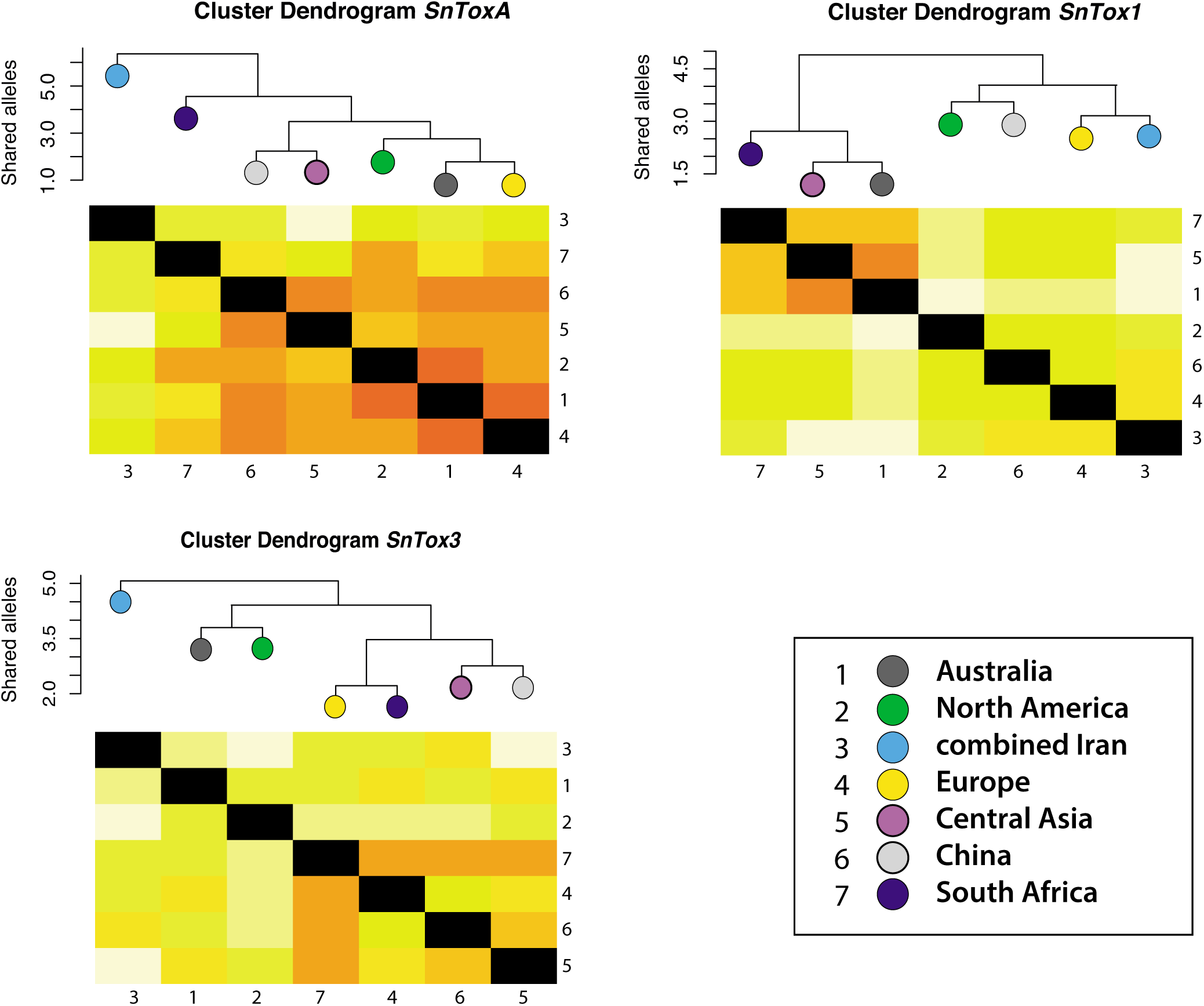
Heatmaps and corresponding cluster dendrograms showing the normalized numbers of shared effector alleles among global populations of *Parastagonospora nodorum*. The lighter the color, the more alleles are shared. The combined Iranian population appears to act as a hub (source population) for globally-shared alleles of *SnToxA* and *SnTox3*, but the pattern is not clear for *SnTox1*.

